# Enhancing the biological activity of polyphenols based on understanding their chemistry

**DOI:** 10.64898/2026.06.01.729321

**Authors:** Ana Bethsy Aguilar-Carrillo, Sofia Antonia Garduño-Valdovinos, Gerardo M. Nava, Vanessa Sánchez-Quezada, Luis Alberto Madrigal-Perez

## Abstract

Polyphenols are compounds synthesized by plants as part of their chemical defense system to counteract biotic and abiotic stressors. These compounds share two key chemical characteristics: their aromatic groups make them insoluble in water, while hydroxy groups provide redox properties. These characteristics may explain how polyphenols interact with mitochondrial membranes (which are lipophilic) and participate in redox (electron scavenging) reactions of the electron transport chain, ultimately affecting ATP synthesis via oxidative phosphorylation. This interaction accounts for both the beneficial and adverse effects of polyphenols. However, no research has examined how hydroxyl groups or a lipophilic environment influence the biological activity of polyphenols. Therefore, this study aimed to explore the impact of hydroxy groups and a lipophilic environment on the biological activity of polyphenols. We tested four polyphenols (quercetin, naringenin, resveratrol, and gallic acid) with varying numbers of hydroxyl and other functional groups to determine how hydroxyl groups affect their biological activity (toxicity) in *Saccharomyces cerevisiae*. Additionally, we evaluated different fatty acids to understand how a lipophilic environment influences polyphenol biological activity. The results of this study support the two main ideas of our hypothesis: 1) a lipid solvent increases the toxicity of polyphenols, and 2) the molecule with the most hydroxyl groups is the most toxic (as seen with quercetin, which has five hydroxyl groups). Consequently, the increased toxicity of polyphenols in lipid solvents, along with their association with oxidizable groups, opens the door to the development of new technologies based on polyphenols.

## Introduction

Polyphenols are chemical compounds produced by plants in response to a biotic or abiotic stress [1]. Polyphenols form part of a chemical defense system that counteracts biological attacks [2] or other types of stress, such as mechanical or UV stress. Additionally, polyphenols are associated with health benefits in mammalian models and humans [3]. However, there is intense debate about its pharmacological use due to its small clinical effect, attributed to poor bioavailability [4]. Given the chemical structure of polyphenols, we can enhance their biological activity. A typical structure of these phytochemicals consists of a phenol group, formed from a benzene ring with a hydroxy group. The benzene ring makes polyphenols poorly soluble in water, and the hydroxy group is highly reactive in redox reactions [5]. These chemical characteristics explain the inhibition of oxidative phosphorylation as the primary target of polyphenols such as resveratrol [6] and quercetin [7]. These molecules can interact with the inner mitochondrial membrane (a lipophilic environment) [8, 9], and the redox potential allows electrons to be removed from the electron transport chain [10]. In turn, inhibition of oxidative phosphorylation accounts for the main biological activities attributed to polyphenols, including antibiotic, anticancer, and antioxidant activities, among others. Also, the low solubility of polyphenols in water may be a critical characteristic that helps explain their apparent low bioavailability. Thus, it is expected that the interaction of polyphenols with lipophilic matrices prompts their biological activity. Moreover, the evidence indicates that electron scavenging by polyphenols in the electron transport chain is responsible for diminishing biological activity, and the functional group responsible for this is the hydroxy group. Therefore, polyphenols with more hydroxy groups would have a higher biological activity. However, there are no reports on the role of solubility or hydroxyl groups in the biological activity of polyphenols. Thus, this study aimed to elucidate the biological role of lipophilicity and hydroxyl groups of polyphenols. Herein, we found that hydroxy groups are essential for UV absorption, and that fatty acids improve the cytotoxicity of polyphenols in *Saccharomyces cerevisiae*.

## Material and methods

### Strain

Experiments were performed using the *S. cerevisiae* BY4742 genetic background (Matα; *his3*Δ*1; leu2*Δ*0; lys2*Δ*0; ura3*Δ*0*) obtained from EUROSCARF (Frankfurt, Germany).

### Culture media

*S. cerevisiae* was activated by taking an aliquot from a vial stored at -20°C and then maintaining it in YPD medium. YPD medium was prepared with 1% yeast extract (BD Bioxon, CDMX, Mexico), 2% casein peptone (BD Bioxon), and 2% glucose (Meyer, CDMX, Mexico). For the experimentation, a complete synthetic medium (SC) was used, which was formulated with 0.18% nitrogen base for yeasts without amino acids (Sigma-Aldrich, Darmstadt, Germany), 0.5% ammonium sulfate (J.T. Baker, Phillipsburg, NJ, USA), 0.2% dipotassium phosphate (J.T. Baker), and 1% dropout mixture without uracil (Sigma-Aldrich). It was supplemented with 400 μg/mL uracil (Sigma-Aldrich) and two concentrations of glucose (0.1% or 5%).

### UV absorption assay

UV absorption was performed with ethanol polyphenol solutions at 100 μM, verting 3 μL onto a μDrop plate. Then, spectral absorbance was measured using the Multiskan Sky instrument (Thermo-Scientific, Waltham, MA, USA), recording absorbance every 2 nm from 200 nm to 500 nm.

### Cytotoxicity test

Cytotoxicity was measured with the exponential growth through the specific growth rate. Growth curves were used to obtain the kinetic parameters as follows. Yeast was grown in 3 mL of YPD medium supplemented with 2% glucose overnight with constant shaking at 30°C. Subsequently, 10 μL of the cell suspension and 137 μL of SC medium supplemented with three levels of quercetin, naringenin, gallic acid, and/or resveratrol (0.1 μM, 100 μM, or 1000 μM) in combination with two concentrations of glucose (0.1 or 5%) were placed in a 96-well microplate. 3 μL of ethanol polyphenol solutions were put into the plate, and as a vehicle control, 3 μL of absolute ethanol was used. The plate was incubated at 30°C with constant shaking for 24 h using the Multiskan Sky instrument (Thermoscientific, Waltham, MA, USA), and absorbance readings at 600 nm were taken every 30 min. The specific growth rate was calculated using the exponential growth equation and fitted in GraphPad Prism for macOS (v. 10.6.1).

### Lipid matrix toxicity assays

Assays to evaluate the effects of a lipid matrix on the cytotoxicity of polyphenols were performed in the same manner as the cytotoxicity assay, except that three fatty acids with different levels of saturation were added to the experimental design: linoleic acid (18:2), oleic acid (18:1), and palmitic acid (16:0), which correspond to a polyunsaturated, monounsaturated, and saturated fatty acid, respectively. In the experimental design, fatty acids were supplemented from an ethanol solution to obtain a final concentration of 1 mM, with absolute ethanol as the vehicle control without fatty acids. Then, in a microplate (96 wells) 10 μL of the cell suspension and 137 μL of SC medium supplemented with 1.5 μL of quercetin, naringenin, gallic acid or resveratrol to reach concentration of 0.1 μM, 100 μM, or 1000 μM in combination with two concentrations of glucose (0.1% or 5%) and 1.5 μL of fatty acids were placed. The plate was incubated at 30°C with constant shaking for 24 hours using the Multiskan Sky instrument (Thermo Scientific, Waltham, MA, USA). Absorbance readings at 600 nm were taken every 30 minutes. The specific growth rate was calculated using the exponential growth equation and analyzed with GraphPad Prism for macOS (version 10.6.1).

### Viability percentage

The specific growth rate, calculated from the exponential phase, represents an indirect parameter of cell division. For this reason, the specific growth rate was used to calculate the viability percentage, obtained as follows: dividing the specific growth rate of the supplementation assay by the mean of the control, then multiplying by 100 to obtain the viability percentage. Thus, control represents 100% of the viability percentage since they are divided by their own values.

### Statistical analysis

The assays were performed in at least three independent experiments, with technical duplicates. All statistical analyses were performed using GraphPad Prism for macOS (version 10.6.1).

## Results

### Selecting polyphenols to test the hydroxy group hypothesis of biological activity

This study aimed to determine whether polyphenol hydroxy groups are responsible for chemical activities, such as cytotoxicity and UV protection. To test this idea, we chose according to chemical characteristics four different polyphenols: quercetin, naringenin, resveratrol, and gallic acid (**Fig. 1**). Quercetin is composed of two benzene rings connected by a heterocyclic pyrone ring, with five hydroxy groups, has a double bond and a carbonyl group (**Fig. 1**). Naringenin has the same chemical characteristics as quercetin, but with only three hydroxyl groups, which is a crucial characteristic to discard the role of the phenol groups in biological activity (**Fig. 1**). Resveratrol is a non-flavonoid polyphenol with a stilbene core structure, composed of two phenolic rings connected by an ethylene bridge with three hydroxy groups and any additional functional group (**Fig. 1**). Gallic acid is a 2,3,4-trihydroxybenzoic featuring both hydroxyl (three) and carboxylic acid (one) functional groups (**Fig. 1**). Those polyphenols allow us to test the importance of hydroxy groups and the role of other functional groups in polyphenols’ main chemical functions.

**Fig. 1.**
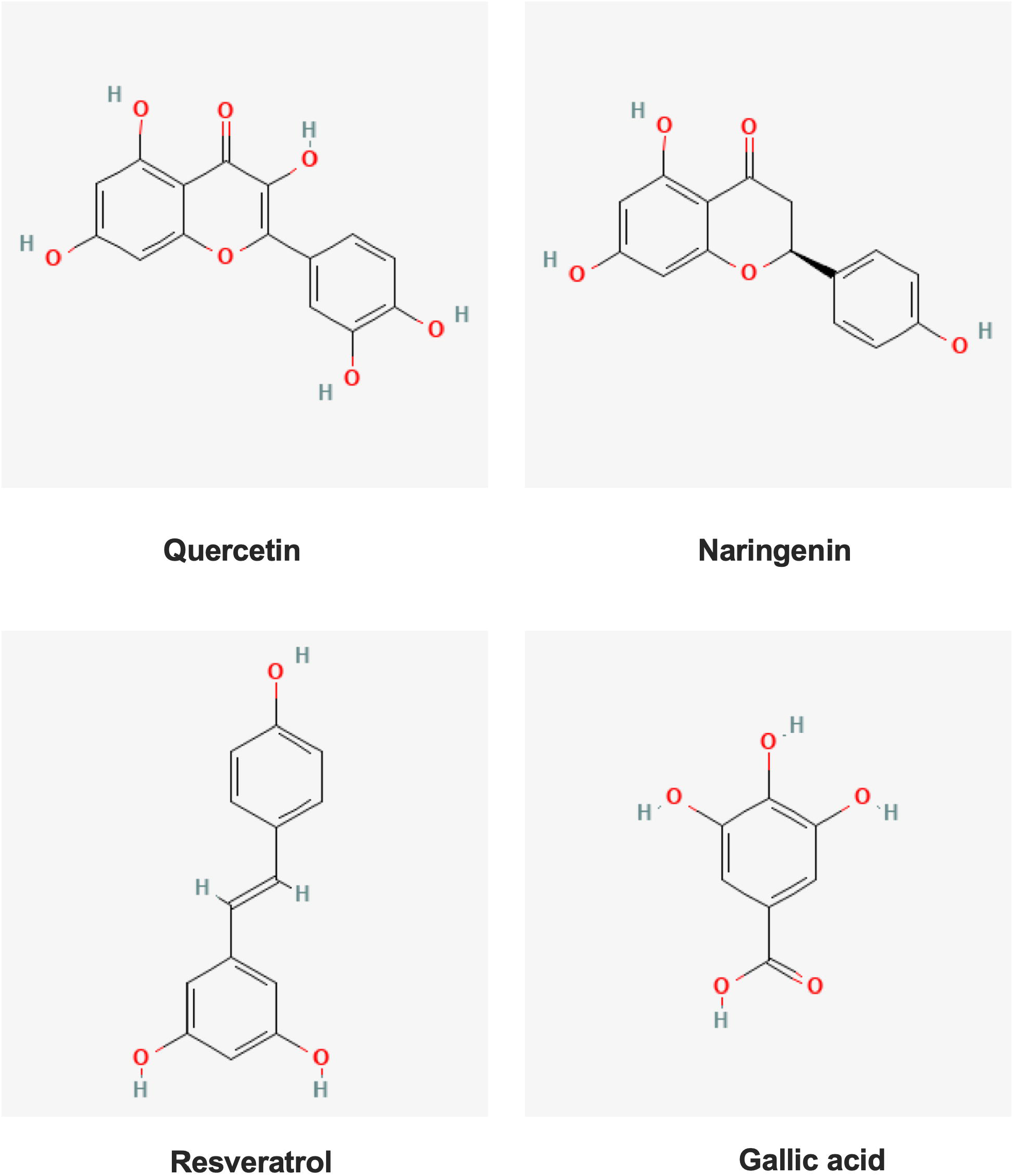
Polyphenol molecular structure. Molecular representations of the polyphenols utilized for this study.

### Association of polyphenol molecules with UV absorption

The first test to prove our hypothesis was to measure the UV absorption of the polyphenols, one of the outstanding functions of these phytochemicals. Also, UV absorption is a non-biological activity that could help determine whether there is a relationship between physical and biological activities of polyphenols. Quercetin showed the highest UV absorption, followed by gallic acid, resveratrol, and finally naringenin (**Fig. 2**). This first test indicates that the number of hydroxy groups is the relevant chemical feature for UV absorption.

**Fig. 2.**
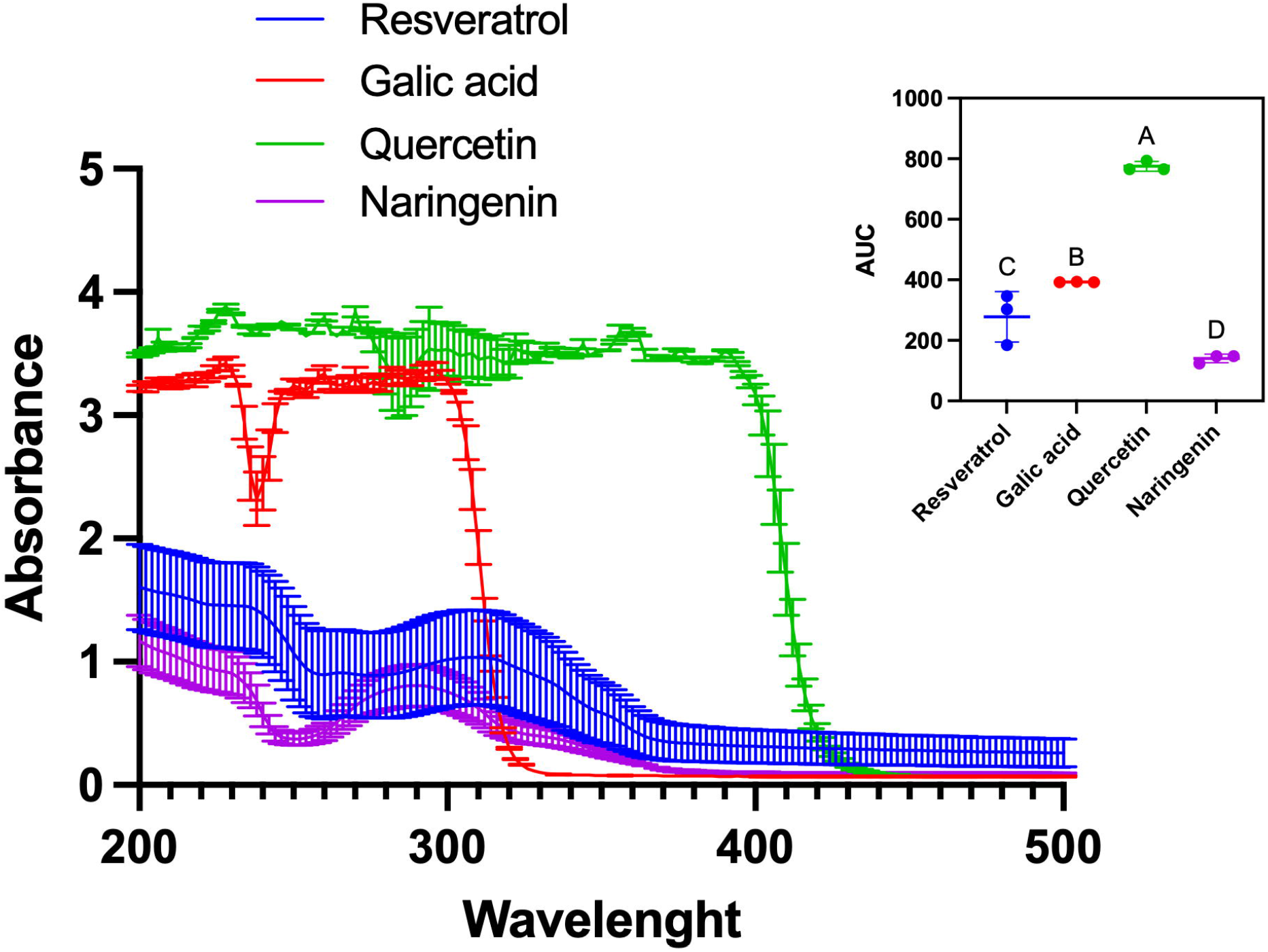
UV absorbance of quercetin, naringenin, resveratrol, and gallic acid. UV absorbance of polyphenols was measured at 100 μM of each polyphenol. The data in the small graph represent the area under the curve (AUC) analysis, with results reported as mean values ± standard deviation from three independent experiments. Statistical analyses were performed using one-way ANOVA followed by the Tukey test; different letters indicate statistical differences.

### Influence of hydroxy groups in polyphenols’ toxicity in S. cerevisiae

Based on the evidence that polyphenols’ main target is the inhibition of ATP synthesis via oxidative phosphorylation [11-13]. Previously, we established a model in *S. cerevisiae* to evaluate the metabolic effects of polyphenols by leveraging its capacity to shift its energetic metabolism in response to glucose concentration: under low glucose concentrations, it primarily uses mitochondrial respiration, and under high glucose concentrations, it primarily uses fermentation [14], this model allow us to deciphering that polyphenols are more cytotoxic in low glucose concentration and led us to propose that main target is the inhibition of ATP synthesis via oxidative phosphorylation [6, 7]. For these reasons, we suggest using polyphenols’ cytotoxicity as an indicator of biological response to elucidate the roles of hydroxyl groups and lipophilic solvents. Thus, under this model, quercetin, resveratrol, naringenin, and gallic acid affect the exponential growth of *S. cerevisiae* in a similar pattern to that of UV absorption (**Fig. 3**). Quercetin and naringenin had opposite effects on exponential growth; quercetin had the most adverse effect, while naringenin had a mild impact (**Fig. 3**). This is important because quercetin chemically only has two additional hydroxy groups than naringenin (**Fig. 1**). Therefore, these data suggest that the number of hydroxy groups in polyphenols is essential for UV absorption and is also related with its cytotoxicity to *S. cerevisiae*.

**Fig. 3.**
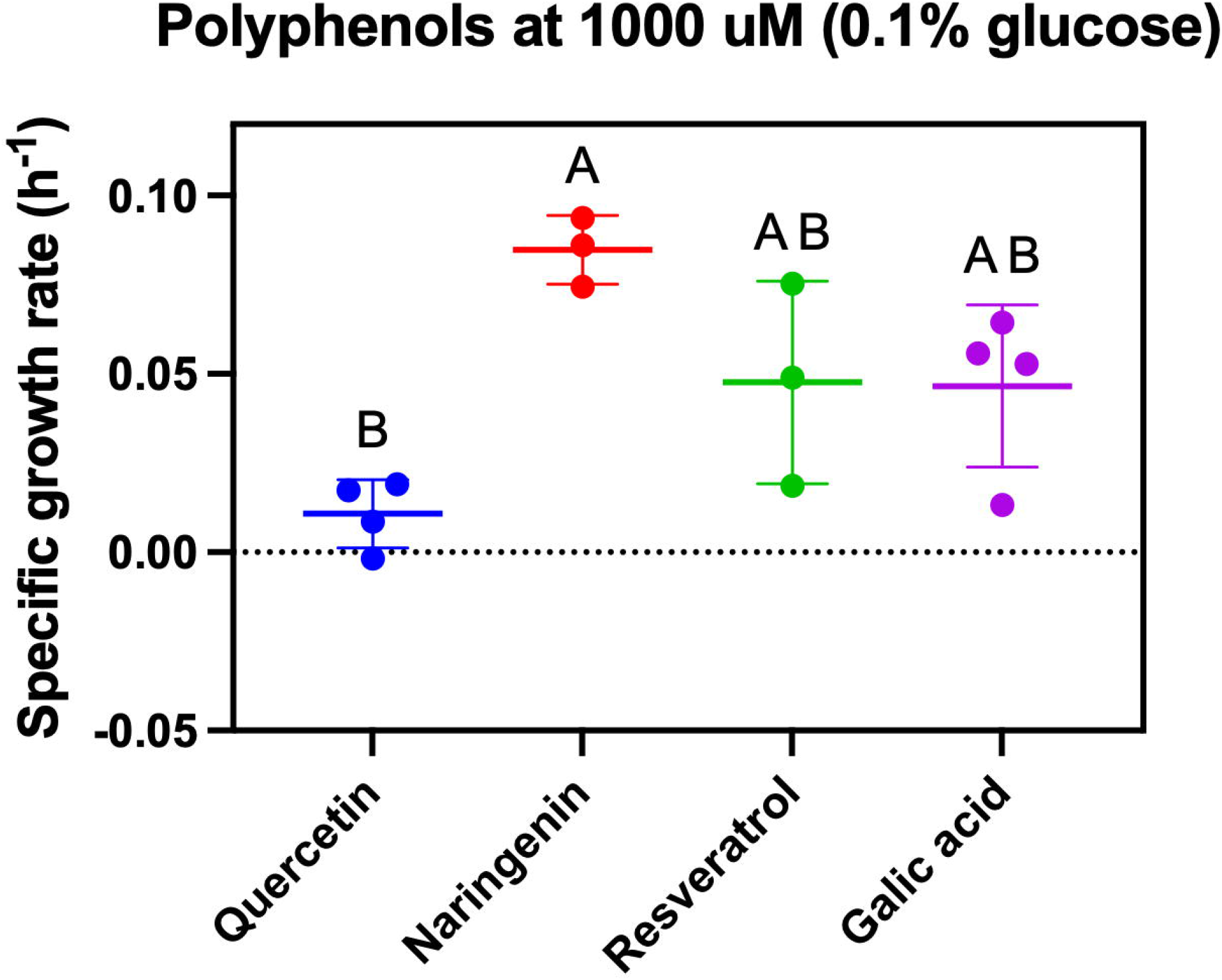
Quercetin, naringenin, resveratrol, and gallic acid affect the exponential growth rate of *S. cerevisiae* at 1000 μM. The effect of polyphenols at 1000 μM was assessed in the presence of 0.1% glucose. The results represent mean values ± standard deviations from three to four independent experiments, including two technical repetitions. Statistical analyses were performed using one-way ANOVA followed by the Tukey test; different letters indicate statistical differences.

### Effect of fatty acid on cytotoxicity of polyphenols in S. cerevisiae

Polyphenols’ low water solubility is expected given their chemical structure, which consists of benzene rings. Based on this, we propose that, in addition to hydroxy groups, the solubility of polyphenols in a lipophilic matrix could enhance their biological activity. To test this idea, we supplemented the culture media with three fatty acids—linoleic acid (18:2), oleic acid (18:1), and palmitic acid (16:0)—representing polyunsaturated, monounsaturated, and saturated fatty acids, respectively. Fatty acids were supplemented to the media culture at 1 mM. Also, for this experimental design, two glucose concentrations were employed: 0.1% glucose, which promotes respiratory growth, and 5% glucose, which mainly supports fermentative growth. This is necessary because polyphenols affect oxidative phosphorylation, and it is required to account for the effects of glucose repression in our model *S. cerevisiae*.

First, we found that quercetin cytotoxicity was augmented with unsaturated fatty acids: oleic acid and linoleic acid at 0.1% glucose (**Fig. 4a-b**), and a pronounced effect was found at 5% glucose with oleic acid (**Fig. 4c**). Cytotocity was also increased with naringenin and oleic acid at 5% glucose (**Fig. 5b**), since naringenin is chemical analog to quercetin it indicates that oleic acid is a suitable solvent for those chemical structures making more bioavailable these phytochemicals. However, no effect on cellular viability was observed at 0.1% glucose when naringenin was supplemented (**Fig. 5a**). In the case of resveratrol, a mixed effect was observed at 0.1% glucose an increased cytotoxicity was observed when resveratrol was supplemented with oleic acid (**Fig. 6a**), while at 5% glucose with linoleic acid (**Fig. 6b**). Finally, gallic acid showed a drastic decrease in viability percentage at 0.1% glucose when was supplemented with palmitic acid, the more likely structure to this polyphenol among fatty acids tested (**Fig. 7a**); and any effect was observed with the other fatty acids or at 5% glucose (**Fig. 7b**). Second, it was also evident that glucose concentration impacts the cytotoxicity of polyphenols. According to reports, we expected polyphenol toxicity to be more pronounced at 0.1% glucose, as this is the glucose concentration at which mitochondrial respiration is more active [15]. However, a mixed effect was found. Quercetin and naringenin exhibited greater toxicity with oleic acid supplementation and 5% glucose (**Fig. 4-5**). On the other hand, resveratrol (supplemented with oleic acid) and gallic acid (supplemented with palmitic acid) showed a marked cytotoxicity at 0.5% glucose (**Fig. 6-7**).

**Fig. 4.**
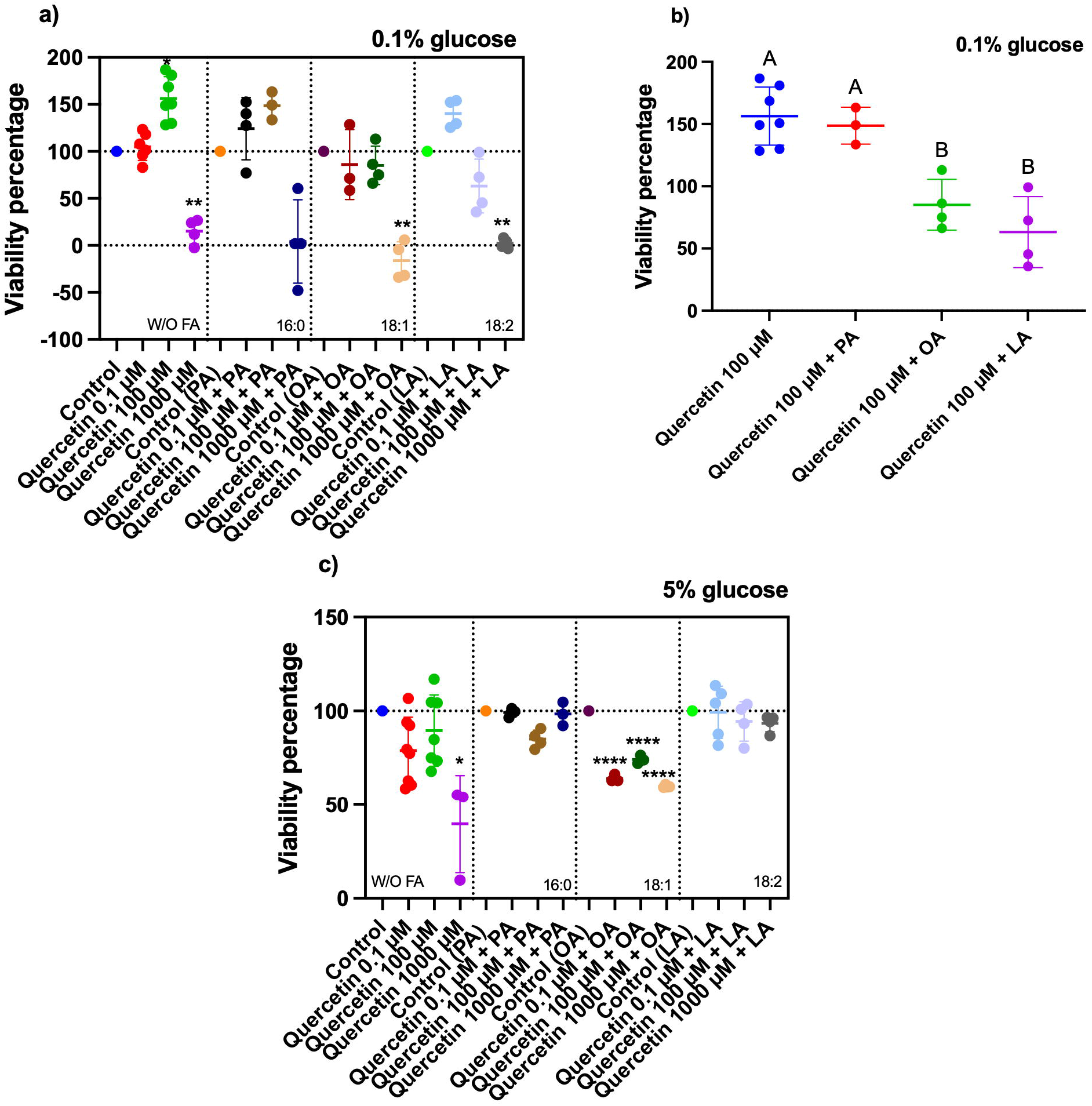
Influence of quercetin and fatty acid supplementation on *S. cerevisiae* cellular viability. Three fatty acids were utilized to study their effect on cellular viability in the presence of quercetin. The three fatty acids used were linoleic acid (18:2; LA), oleic acid (18:1; OA), and palmitic acid (16:0; PA). Viability percentage of *S. cerevisiae* at 0.1% glucose. b) Comparison of the effect of the different fatty acids with quercetin at 100 μM. c) Viability percentage of *S. cerevisiae* at 5% glucose. The results represent mean values ± standard deviations from 3 to 8 independent experiments, with two technical repetitions. For panels a) and c), statistical analyses were performed using one-way ANOVA followed by the Dunnett test vs. each control (*\*P* ≤0.05; *\*\*P*≤0.01; *\*\*\*\*P*≤0.0001). Panel b) statistical analyses were performed using one-way ANOVA followed by the Tukey test; different letters indicate statistical differences.

**Fig. 5.**
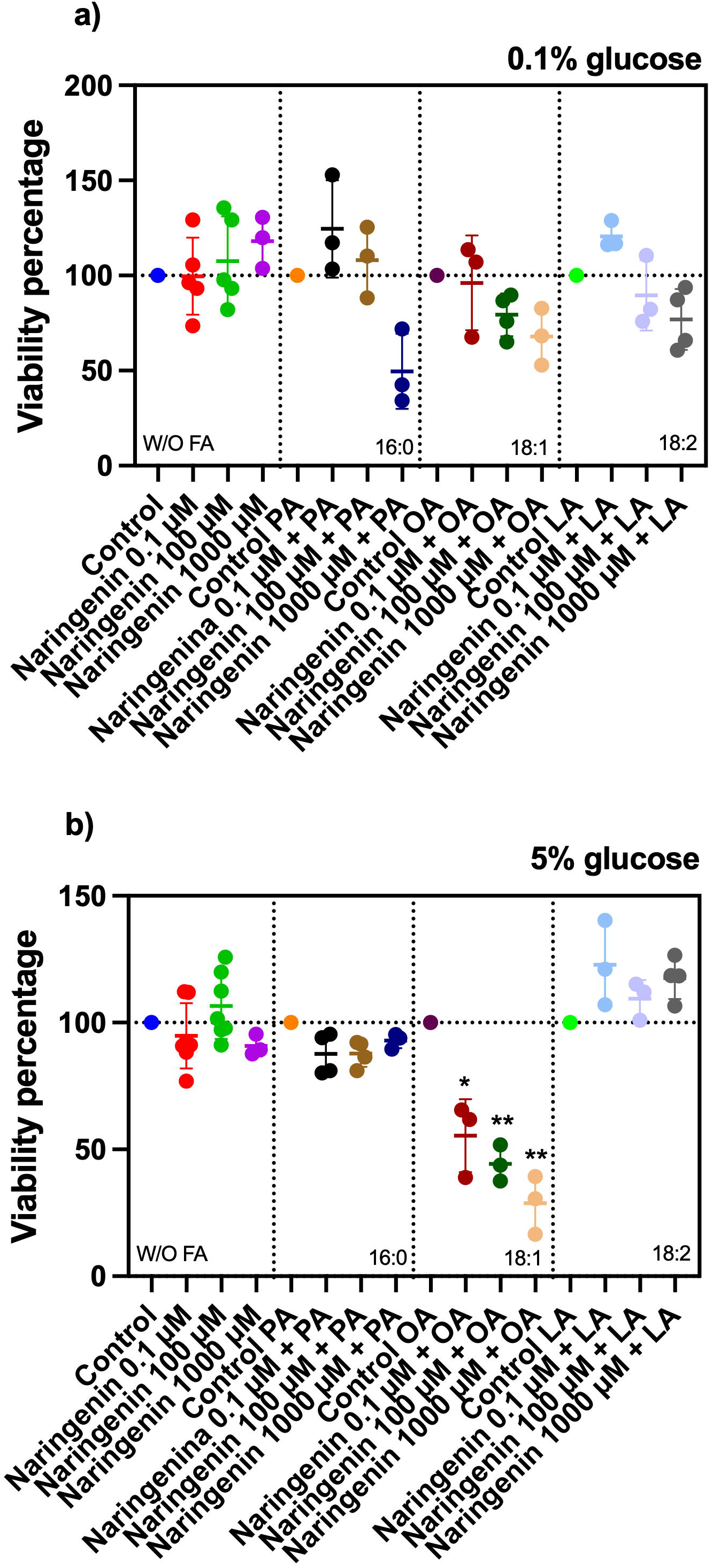
Naringenin and fatty acid supplementation effect on *S. cerevisiae* cellular viability. Three fatty acids were used to investigate their impact on cellular viability in the presence of naringenin. The fatty acids examined were linoleic acid (18:2; LA), oleic acid (18:1; OA), and palmitic acid (16:0; PA). a) Viability percentage of *S. cerevisiae* at 0.1% glucose. b) Viability percentage of *S. cerevisiae* at 5% glucose. The results represent mean values ± standard deviations from 3 to 6 independent experiments, with two technical repetitions. Statistical analyses were performed using one-way ANOVA followed by the Dunnett test vs. each control (*\*P* ≤0.05; *\*\*P*≤0.01).

**Fig. 6.**
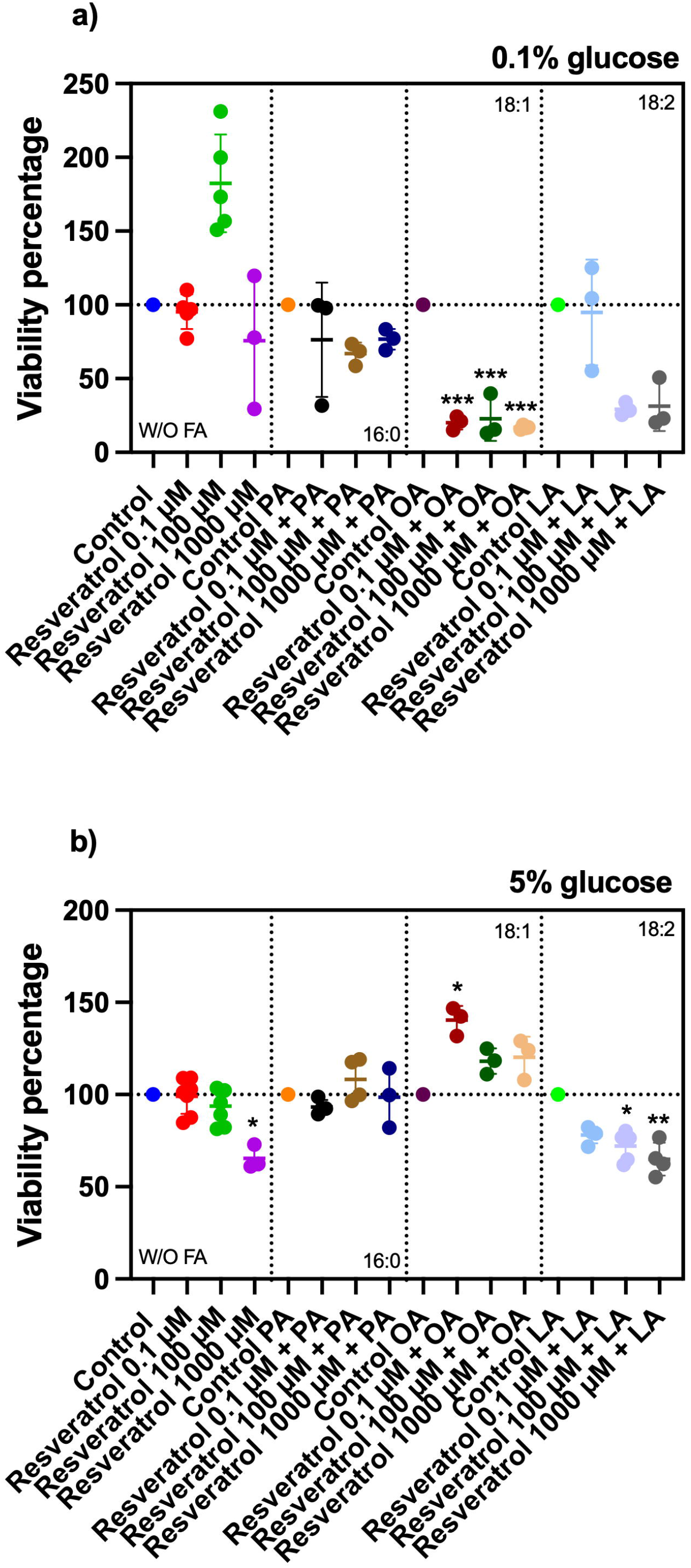
Influence of resveratrol and fatty acid supplementation on *S. cerevisiae* cellular viability. Three fatty acids were utilized to study their effect on cellular viability in the presence of resveratrol. The three fatty acids used were linoleic acid (18:2; LA), oleic acid (18:1; OA), and palmitic acid (16:0; PA). a) Viability percentage of *S. cerevisiae* at 0.1% glucose. b) Viability percentage of *S. cerevisiae* at 5% glucose. The results represent mean values ± standard deviations from 3 to 6 independent experiments, with two technical repetitions. Statistical analyses were performed using one-way ANOVA followed by the Dunnett test vs. each control (*\*P* ≤0.05; *\*\*P*≤0.01;*\*\*\*P*≤0.001).

**Fig. 7.**
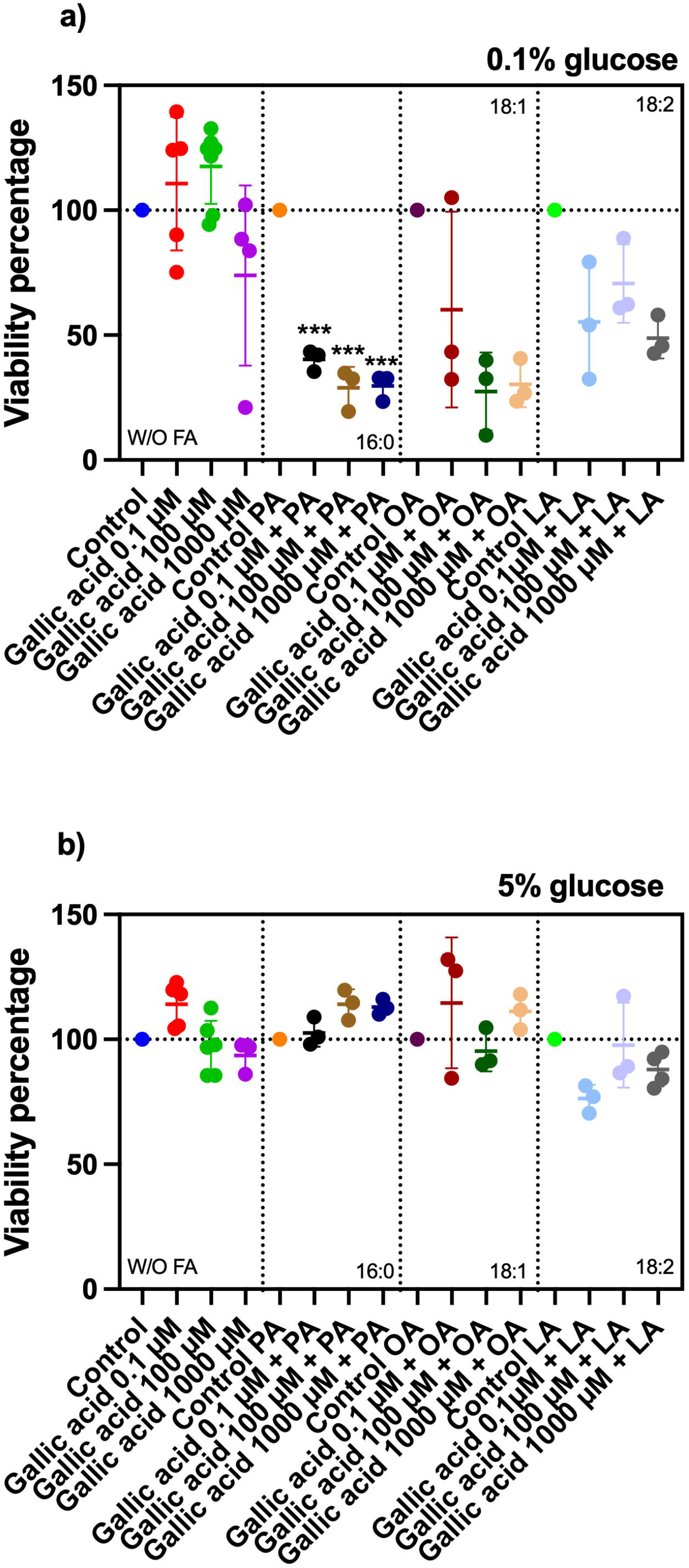
Gallic acid and fatty acid supplementation effect on *S. cerevisiae* cellular viability. Three fatty acids were used to investigate their impact on cellular viability in the presence of gallic acid. The fatty acids examined were linoleic acid (18:2; LA), oleic acid (18:1; OA), and palmitic acid (16:0; PA). a) Viability percentage of *S. cerevisiae* at 0.1% glucose. b) Viability percentage of *S. cerevisiae* at 5% glucose. The results represent mean values ± standard deviations from 3 to 6 independent experiments, with two technical repetitions. Statistical analyses were performed using one-way ANOVA followed by the Dunnett test vs. each control (*\*\*\*P*≤0.001).

Overall, these data indicate that fatty acid supplementation increases polyphenol cytotoxicity (consistent with their chemical similarity) in a glucose-independent manner.

### Association of hydroxy groups, cytotoxicity, and fatty acids

Finally, to gain a better understanding of whether hydroxy groups are the functional groups are responsible for the cytotoxic effect of polyphenols, we performed a multivariate PCA. In this analysis, we incorporated the following continuous variables: number of hydroxy groups, polyphenol concentration, other functional groups (carbonyl and carboxyl), fatty acid saturation level, glucose concentration, and exponential growth. As expected, glucose concentration drives exponential growth, reflecting their vectors in the same quadrant (**Fig. 8**). In contrast, the number of hydroxy groups and polyphenol concentration push the data in the opposite quadrant of glucose concentration and exponential growth demonstrating that hydroxy groups as well polyphenol concentration are the primary influencers of cytotoxic effect of these phytochemicals (**Fig. 8**). The carbonyl group, carboxyl group and fatty acid saturation showed a middle impact according to its vectors to the exponential growth (**Fig. 8**). It was expected for fatty acid instauration since it enhances cytotoxicity of polyphenols according to the chemical analogy and not as a linear effect. Altogether, this study demonstrates that hydroxy groups are the preponderant chemical feature of polyphenols for this chemical activity.

**Fig. 8.**
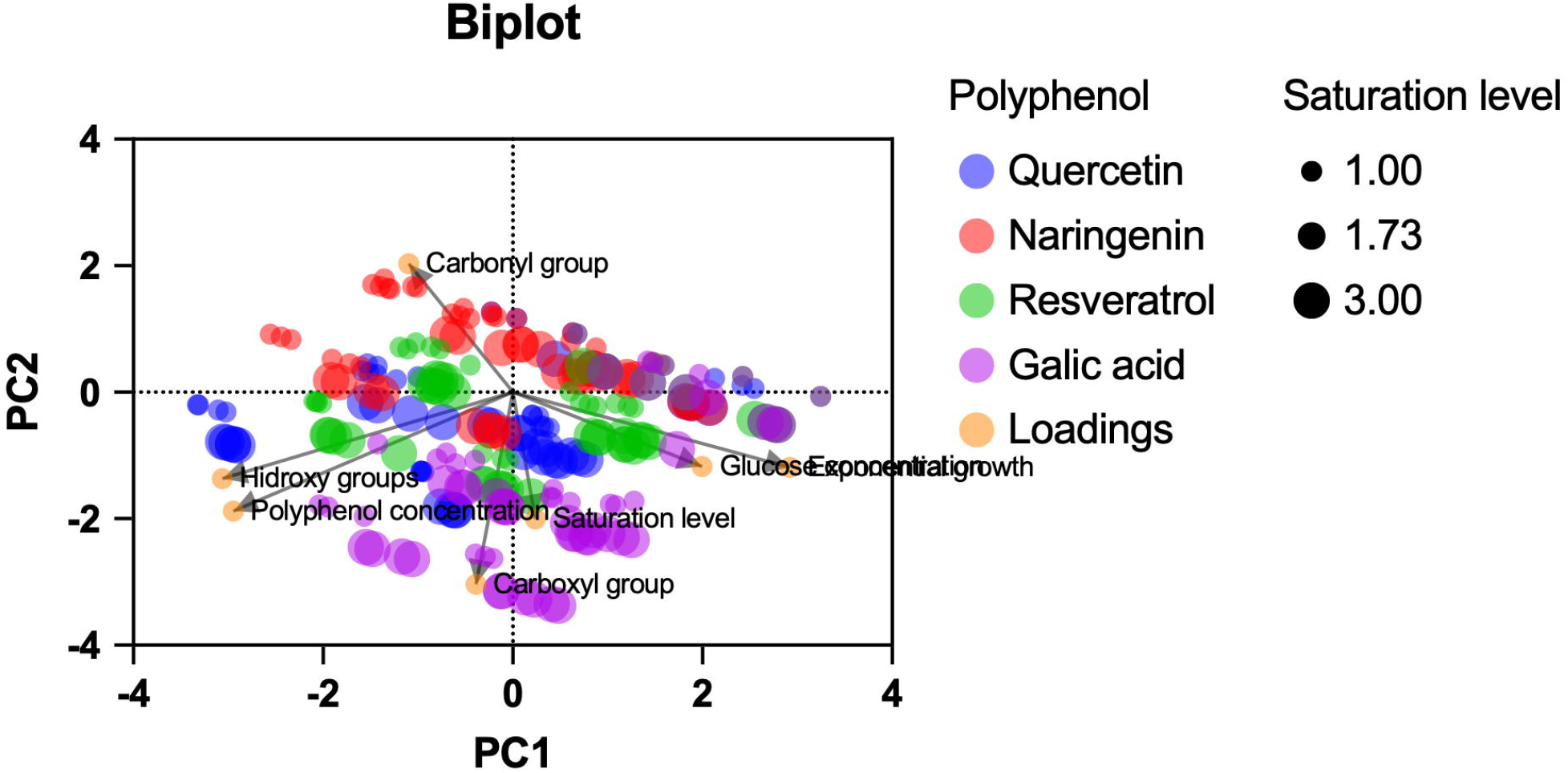
Principal component analysis (PCA) of biological effects of quercetin, naringenin, resveratrol, and gallic acid on *S. cerevisiae*. PCA was performed in GraphPad Prism for macOS (version 10.6.1). The method for selecting principal components was based on parallel analysis with Monte Carlo simulation (1000 simulations) and an auto random seed.

## Discussion

Polyphenols are essential compounds synthesized by plants as part of their chemical defense system, effectively countering a range of biotic and abiotic stresses [16]. Polyphenols possess two critical characteristics: their aromatic structure renders them insoluble in water, while their hydroxyl groups impart significant redox properties. The central hypothesis for the molecular mechanism of action of polyphenols posits that they impair oxidative phosphorylation [11, 12]. Accordingly, the chemical attributes of polyphenols explain how they can interact with mitochondrial membranes and scavenge electrons from the electron transport chain, thereby influencing oxidative phosphorylation and producing both beneficial (in mammalian models) and adverse effects. Despite this understanding, research on the impact of hydroxyl groups and the lipophilic environment on the physical or biological activity of polyphenols is currently insufficient. This study decisively addresses this gap by investigating how these factors influence polyphenol activity using the yeast species *S. cerevisiae* as a biological model. We tested four distinct polyphenols (quercetin, naringenin, resveratrol, and gallic acid), each with differing numbers of hydroxyl and functional groups, to rigorously assess the influence of hydroxyl groups on their biological activity. Moreover, we evaluated the polyphenols in conjunction with various fatty acids to determine how a lipophilic environment affects their activity. The results demonstrate that the number of hydroxyl groups is crucial for polyphenol chemical activity (UV absorbance and biological activity), particularly cytotoxicity. Importantly, we found that the biological activity of polyphenols significantly increases in the presence of fatty acids. These findings robustly support our two main hypotheses: 1) a lipid solvent enhances polyphenol toxicity, and 2) the polyphenol with the highest number of hydroxyl groups—quercetin, which has five hydroxyl groups— exhibits the most significant toxicity. Consequently, the observed increase in polyphenol toxicity in lipid solvents, along with their association with oxidizable groups, opens exciting avenues for the development of new technologies that account for these findings.

It is well known that the antioxidant capacity of polyphenolic compounds significantly depends on the number of hydroxyl groups [17]. However, the influence of polyphenol hydroxylation on *in vitro* protein activities has also been reported. For example, the number of hydroxy groups in polyphenols, particularly flavonoids, prompts inhibition of PKCδ phosphorylation [18]. Moreover, hydroxylation in the B rings of polyphenols affected the VEGF-inhibitory activity; the potency order of the B-ring series of flavonols, from highest to lowest, was: myricetin (3,5,7,3’,4’,5’), quercetin (3,5,7,3’,4’), kaempferol (3,5,7,4’), and galangin (3,5,7) [19]. Additionally, a strong positive correlation (R^2^ = 0.9873) was observed between the total number of hydroxyl substitutions and the potency of flavonoids in inhibiting VEGF-induced activation of VEGFR-2 [19]. Nonetheless, no study has examined how hydroxylation affects the biological activity of polyphenols. As expected, quercetin, the polyphenol with the highest number of hydroxy groups, showed the most significant cytotoxic phenotype (**Fig. 3**). On the contrary, naringenin, the same molecule as quercetin but without the two hydroxy groups in the B ring, displayed the lowest cytotoxic effect (**Fig. 3**). Overall, these data indicate that hydroxylation is most influential chemical characteristic of polyphenols for its activity.

The hydrophobicity of polyphenols is a well-described property of these molecules, evident by their poor solubility in water. Also, the interaction of polyphenols with biomembranes is widely described. For example, the localization of resveratrol within the membrane bilayer affects its fluidity in a dose-dependent manner, correlating with membrane rigidity and the cytotoxic effects of resveratrol on the MCF-7 breast cancer cell line [8]. Additionally, the detoxification of resveratrol in the liver and intestine in humans primarily involves phase II enzymes, which enhance its solubility by adding sulfate and glucuronide groups [20]. Another example is quercetin, which has been proposed as a potential coenzyme Q mimetic molecule, with its primary action occurring in the mitochondria [21]. Research has shown that SH-SY5Y cells can accumulate quercetin (10 μM) in their mitochondria [22]. Furthermore, the amphipathic properties of quercetin enable it to modify cell membrane permeability. It has been reported that when quercetin is encapsulated in liposomes (specifically Lipoid E80), there are changes in the structural properties of the liposomes. This includes a reduction in membrane thickness due to the intercalation of quercetin between the hydrophobic core and the polar heads of the lipids [23]. Despite these reports, there is a lack of investigation into enhancing polyphenol activity through hydrophobicity. In a previous report, our research group found that quercetin enhances its cytotoxic properties when supplemented with fatty acids [24]. This observation prompted us to test fatty acids and to associate them with the chemical structure of polyphenols. Interestingly, we found that the cytotoxicity of all the polyphenols tested was augmented in the presence of fatty acids (**Fig. 4-7**). There is no association between fatty acid instaurations and this increase in cytotoxicity in polyphenols (**Fig. 8**), which is more related to the chemical similarity between polyphenols and fatty acids. Altogether, these findings strongly suggest that polyphenol properties might be enhanced when solubilized in hydrophobic matrices.

Our findings clearly establish two key points: 1) lipid solvents enhance polyphenol toxicity, and 2) quercetin, with its five hydroxyl groups, exhibits the highest toxicity. This increased toxicity in lipid solvents, linked to oxidizable groups, paves the way for innovative technological advancements.

## Acknowledgments

This study was funded by grants from Tecnológico Nacional de México (20026.24-PD) and PICIR 2022 program from the Instituto de Ciencia, Tecnología e Innovación del Estado de Michoacán de Ocampo (Grant: PICIR22-008-C).

## Data availability statement

The data that support the findings of this study are available from the corresponding author upon reasonable request.

## Author contribution

All authors contributed to the study conception and design. Material preparation, data collection, and analysis were performed by Ana Bethsy Aguilar-Carrillo and Sofia Antonia Garduño-Valdovinos. The first draft of the manuscript was written by Luis Alberto Madrigal-Perez and all authors commented on previous versions of the manuscript. All authors read and approved the final manuscript.

